# Workflow for Protein N-terminal Acetylation and C-terminal Amidation Application using CcpNmr Analysis v2.4 Platform and Aria2.3 Structure Calculation Software

**DOI:** 10.1101/2020.12.08.414565

**Authors:** Bruno de Paula Oliveira Santos, Bruno Marques Silva, Mariana Torquato Quezado de Magalhães

**Affiliations:** Programa Interunidades de Pós-Graduação em Bioinformática, Universidade Federal de Minas Gerais, Av. Antonio Carlos, 6627, Belo Horizonte, Minas Gerais, 31270901, Brazil; Departamento de Bioquímica e Imunologia, Universidade Federal de Minas Gerais, Av. Antonio Carlos, 6627, Belo Horizonte, Minas Gerais, 31270901, Brazil

**Keywords:** workflows, NMR spectrocopy, experimental protocol, characterization, L-phenylseptin

## Abstract

Structural biology is a field that enables a better understanding of proteins from scratch. From the available techniques, solution NMR is one well established that provides structure, dynamics and protein-molecules interaction. In a NMR lab routine, from data acquisition until protein/mechanisms elucidation comes a process that can undergo months. During the past decades, different tools were developed for NMR data processing, peaks assignment, structure elucidation and data submission. Since many of these programs demand great computational skills, a few groups have tried to combine those programs and make them more friendly and useful, what can possibilite a faster process. Here we highlight CCPNMR2.4 analysis and ARIA2.3, responsible for peak assignment and structure calculation, respectively, and can work associated. Although being academic free and the possibility of working with a GUI interface, the common N-terminal acetylation and C-terminal amidation modifications are not implemented in a way that possibilities to work with them in combination, what results in a dilemma. This work brings visual data that evidences the low usability of CCPN and ARIA with N-terminal acetylated and C-terminal amidated proteins and propose a workflow to overcome this problem, which may improve the usage of both software in the mentioned versions and facilitate the lab users already used to these programs. As a proof of concept, we have chosen a N-terminal amidated peptide, L-Phenylseptin, whose structure has already been solved with other programs. Statistical data shows that no significant difference was found with the structure obtained with the new protocol. In conclusion, we exhibit a new protocol that can be used in combination with CCPNMR2.4 and ARIA2.3 for protein with the mentioned modifications and it successfully works and manipulates these molecules.

## 1. Introduction

Applications for NMR structural biologists have been in development inside big NMR groups according to their necessities. Starting from assignments in papers and migrating to computer interface spectra interactions, NMR software have grown in usage and tools for researchers since the end of the 80’s. If the established software do not reach your needs, why not develop your own system? Quality is synonym of improved producing, results and also, better work. When your group is able to develop your own software, features as usability can be implemented considering their own tastes and new implementations can go according to predefined models. Sometimes, the available platforms already accomplish the groups needs and just need some adaptations, what, in time, results in new versions of the same software. However, scientific computation is not an easy task, and multi-collaborative groups are set up to integrate NMR data and software development.

Historically, NMR software for protein assignment was first developed in 1989, when Ansig was published and made available (Kraulis 1989). Kraulis describes the laborious job of peak assignment in papers and how an interface guided program written in Fortran 77 could make the process faster and more practical, allowing the user to check the lists before exporting them for the structure software (in the work, X-PLOR was cited). In 1991, EASY was published as a new software for 2D spectra interactive assignment (Eccles et al. 1991) that was quickly substituted in 1995 by XEASY, an improved version that contented the assignment of 3D and 4D spectra and with the concept of ‘strips’, to visualize 3D spectra in a 2D manner (Bartels et al. 1995). Johnson and Blevins, in 1994, worked in NMR analysis integration with the structural data and also, integrate multiple spectra, instead of working with them “one at a time”, originating the NMR View (Johnson and Blevins 1994). Sparky, from UCSF, had its first published version in 1999, first released for Linux systems (Goddard and Kneller 1999), and the content was similar what the other ones already provided: 2D and 3D NMR analysis and assignment. In 2007, Burrow-owl software was created and its procedure relays on transposing the NMR spectra, projecting a slice, combination of two spectra, diagonal projection and integration of the spectra in an one fewer order dimension (Benison et al. 2007). Nowadays, Ansig was substituted by the collaborative platform CcpNmr Analysis, NMR View was upgraded for NMRFx and SPARKY maintenance was ceased in 2001, being substituted in 2015 by NMRFAM Sparky (Vranken et al. 2005; Norris et al. 2016; Lee et al. 2015). They work in most known and used operating systems (Linux, Windows and MacOs) and are supported in more recent script languages, such as Python and Java.

Even before the Ansig development, a software for protein structure calculation was already available in 1987, the first release of X-PLOR (Brünger 1993). X-PLOR (1.0) had evolved from the CHARMM force field program, and was capable of using X-ray crystallographic and NMR data for determine structural models, using molecular dynamics for structure refination. In 1998, the same author of X-PLOR and new collaborators created a new definition of structure calculation software, the CNS. Using an own low-level script language, CNS is versatile in the source experiments, being used solution NMR, X-ray crystallography and also for solid state NMR and electron microscopy. CNS, over the X-PLOR 1.0, improved with the possibility of using more computational power and speeding the calculation process (Brünger et al. 1998). Both software are no longer maintained and were incorporated by NIH group, creating a new tool, the Xplor-NIH, which uses a variety of minimizations process to improve structure calculation (Schwieters et al. 2003). In other vertent, Nilges and collaborators in 1997, implemented a workflow for working with ambiguous NMR signs and it used X-PLOR for structure calculation (Nilges et al. 1997). In 2001 the situation changed, when the workflow was released as a software for NOE ambiguities and structural calculation, integrated with CNS, named Aria v1.0. Aria received several updates through the years and is one of the main free academic options for structural calculation nowadays (Linge et al. 2001; Rieping et al. 2007). Cyana is also a main established software and its precursor, CANDID, was developed in 2002, that had as a main new component at the time, the automated NOE assignment (Herrmann et al. 2002). CANDID was reformulated as CYANA in 2004 to improve the handling of NMR chemical shift ambiguities (Güntert 2004).

New advances of 2010 until nowadays includes the outcoming of virtual machines, such as NMRbox, that allows researchers around the world to use a supermachine located in an US cluster. The platform provides pre-installed NMR software that allows the user to skip a dispendious times trying to install each software (Maciejewski et al. 2017). Another interesting outcome is the possibility of making your tasks in an online web server, such as the new version of Aria that, besides the downloadable version, allows the user to use their super machines for structure calculation and provides structure visualization for data evaluation (Allain et al. 2020). It is important to notice, however, that some software lose maintenance since they are created by students during graduate processes, and the research group may lose capacity to keep the development and update of their tools. That is why collaborative projects have become of particular interest in NMR groups. Free academic software are available for each group to use for their own purposes but also allows more people to test and contribute in possible flaws or missing tools, that can be of interest of many other researchers. Here we highlight the importance of two of the quoted software in this process: CcpNmr Analysis v2.4 and Aria 2.3. Like a great union, CcpNMR analysis allows NMR assignment, Aria directly communicates to import the assigned data and make the structure calculation and validation and the results are exported back to Aria for structure analysis (Vranken et al. 2005). For NMR spectroscopists, this combination is very positive and provides a possibility to standardize a method to solve the structure of different proteins.

Here, we report on a workflow that allows to consider the N-terminal acetylation and C-terminal amidation of proteins during structure calculation, using CcpNmr Analysis v2.4 as the assignment platform and Aria2.3 as the structure calculation software. The implementation of these extremities molecules are missing in an integrated manner, what makes CcpNmr does not recognize this information on Aria and vice versa. The N-terminal acetylation and C-terminal amidation interferes in the final structures models and can not lack during the steps of NMR analysis. They guarantee to proteins features as: durability, endurance, binding and recognition. The C-terminal amidation is produced by the action of the peptidylglycine α-amidating monooxygenase (PAM) enzyme when it recognizes dipeptides or tripeptides in proteins sequences, such as GK, GR, GKK, GKR (Prigge et al. 2000). The C-terminal amidation have been found in many bioactive peptides, leading the peptides to have a less ionizable C-terminal, what gives the molecule more chances for executing their biological actions (Dennison and Phoenix 2011; Dennison et al. 2012). Examples of biological activity includes antimicrobial activity, where the C-terminal amidation impairs mechanisms of protein degradation (Mura et al. 2016); neuroactivity, since neuropeptides not amidated are not recognized by nervous systems cells (Eipper et al. 1992); hGLP-1 peptides that are responsible for insulin release and post-prandial plasma glucose lowering (Zhang et al. 2004) and others. Regarding protein acetylation, this modification can occur in the N-terminal portion or at the H∊ of lysine residues. The focus of this study remains on the first option. N-terminal acetylation occurs by a series of acetyltransferases, on translation process of the protein, post-translationally on prokaryotes and on processed regulatory peptides of eukaryotes (Driessen et al. 1985; Dockray 1987). This modification occurs in 85 percent of eukaryotic proteins and in bioactive peptides, such as human defensin-3 (Papanastasiou et al. 2009).

Here we demonstrate a direct and easy-to-implement workflow, that was tested to be used intercalating the already known Aria2 steps for structure calculation. Our observations provides a basis to suggest that this workflow will improve the use of the combination CcpNmr-Aria by researchers that work with post-translational modified proteins and also reduce the use of different software when working with different types of molecules. Aiming to validate this project, we solved the structure of an antimicrobial peptide, the L-phenylseptin. This peptide, from *Hypsiboas punctatus* amphibian, have 14 amino acid residues and is known for its predator warning function, which structure have been previously solved using NmrView and Xplor-NIH and serve as comparison model to the new structures (de Magalhães et al. 2013). Our conclusions suggest that L-phenylseptin final structures are slightly changed from those that are not C-terminal amidated, and highlight the importance of considering these modifications during protein analysis.

## 2. Modifications Implementation

### 2.1. Data Survey

With the purpose of analyse and compare the usage of assignment and structure calculation software, pdb files were collected from Protein Data Bank webserver. The terms used in the search were: a) QUERY: Full Text = “antimicrobial peptide” AND (Experimental Method = “SOLUTION NMR” AND Number of Polymer Residues per Assembly <= 40 AND Polymer Entity Type = “Protein”); b) QUERY: Full Text = “ccpnmr” AND (Experimental Method = “SOLUTION NMR" AND Polymer Entity Type = “Protein”); c) QUERY: Full Text = “ARIA” AND (Experimental Method = “SOLUTION NMR” AND Polymer Entity Type = “Protein”); d) QUERY: Full Text = “xeasy” AND (Experimental Method = “SOLUTION NMR” AND Polymer Entity Type = “Protein”); e) QUERY: Full Text = “sparky” AND (Experimental Method = “SOLUTION NMR” AND Polymer Entity Type = “Protein”); f) QUERY: Full Text = “nmrview” AND (Experimental Method = “SOLUTION NMR” AND Polymer Entity Type = “Protein”). A threshold of 40 amino acids residues was arbitrarily established for the antimicrobial peptides class, to limit the size of these proteins and the quantity of files analyzed. The informations of the purchased files were extracted: a) using command-line Linux tools; b) manually assessed at the PDB webserver, when the “software used” data was not available in the files; c) in the published article, when the option “a” and “b” were not available. When none of the three options were indisposable, the target information was considered unavailable. Graphical visualization was provided by matplotlib library in python 3.6.

### 2.2. NMR Acquisition

This is a subsection (style: IUCr body text)

### 2.3. NMR assignment in CcpNmr Analysis

The first step to add the N-terminal and C-terminal modifications is to insert them in the molecule inside a CcpNmr project. In “add sequence”, the user must put their amino acid residues sequence and at the N-terminal portion the term “Acy” and/or at the C-terminal portion, the term “Nh3”. This extensions come direct in CcpNmr and nothing needs to be installed in this step. In atom browser, the hydrogens atoms are available for the acetyl as “H2*”, and for the amidation, “H” and “H’”. The peak picking process consists of evaluating the spin systems of each residue in intra-residue correlations experiments (e.g. COSY, TOCSY), the inter-residue correlations (e.g. NOESY) and heteronuclear correlations (^13^C/^15^N HSQC) (Sugiki et al. 2017). CcpNmr have combined the DANGLE package that allows the prediction of psi and phi angles. In the “Structure” tab, the user have the “Make distance restraints” option, that converts volume/height NOESY information in atoms distance restraints.

### 2.4. Aria2.3 Project

A new ARIA project is created under the command “aria2 -g” on command line shell. In the tab “CCPN data model”, the path of the CcpNmr project must be inserted. In the CNS tab, the CNS executable path. With the CcpNmr project chosen, the list are recognizable by Aria. The user is then capable of inserting the chemical shift list, molecular system, spectrum, dihedral angles and ambiguous and unambiguous distances directly from the CcpNmr project. Regarding the protein modifications, the user must insert in the tab “Molecular System'': a) Linkage Definition: TOPPALLHDG5.3, b) Topology definition: TOPPALLHDG5.3, c) Parameters Definition: PARALLHDG5.3. Those are the files modified to add the N-terminal acetylation and C-terminal amidation. Is also important to create a new folder to where the calculations will occur: in the tab “Project”, insert the new folder path in “working directory” and “temporary path”. This way, the modification patch will work without further alteration.

To insert the modification, the user must download the zip file (S1) and uncompress it in the same folder where the aria file is located (.xml file). The user must execute the install file, using the following command:

sudo ./install

It is default, after creating the aria project, in the xml extension, to execute the commands:

aria2 -s ariafile.xml

aria2 ariafile.xml

The first command allows ARIA to create the folder where the iterations will occur. The second one is where the ARIA starts the structure calculation. Here, we implement a new step:

aria2 -s ariafile.xml

**./run_patches <structure_folder> <N or C or NC>**

aria2 ariafile.xml

The second command implements the changes in topology and parameters to the new run. The second argument must be the name or path of the folder where the structure runs will occur. The third argument must be N or C or NC, if you want to implement the N-terminal modification or the C-terminal modification or both, respectively. Example:

./run_patches folder01 NC

### 2.5. Files Changes

#### 2.5.1. Aria/CcpNmr

The main strategy to include these modifications was to consider them as residues. That in mind, the “new residues” were included in the files “atomnames.xml” and “iupac.xml”, from aria2.3 folders, with the same atoms nomenclature found in these molecules file of CCPN. They were named “ACY” and “NH2”. The file “AminoAcid.py” was modified to include the acetylation and amidation: The one letter code for both were set as “X”, as already seen in PDB deposited structures. In CcpNmr, the modification was in the folder “molecules”, where the files “others_acy” and “others_nh3” were changed to include a ccp_3_letter code, same as aria2.3 “atomnames” and “iupac” files.

#### 2.5.2. CNS

The main changes are in the CNS files, to include the topologies and parameters. The topology files contain the “new residues” and the molecule bonds. The “topallhdg5.3.pro” has changed the residue ACE to ACY and the atoms nomenclatures. The residue NH2 was included and the “presidue PEPT” was increased with the new bonds and angles to be included in the backbone chain. The parameter file, parallhdg5.3.pro, include the new atoms, bonds, angles, dihedrals and impropers. These parameters were recycled from bonds and atoms with the same features of the amides and acetyl groups. The same was considered when evaluating the dihedral and impropers. (Fig S1).

## 3. Results and Discussion

### 3.1. CcpNmr users tend to choose ARIA during protein solving and vice-versa, but not for N-terminal acetylated and C-terminal amidated proteins

Since the frequency of C-terminal amidation and N-terminal acetylation in antimicrobial peptide is high, because of the importance of these modifications on the functionality of these type of molecules, the antimicrobial peptides N-terminal acetylated and/or C-terminal amidated with ≤ 40 residues were collected from PDB database to evaluate the most frequent software used for assignment and structure calculation (Table 2). Regarding the assignment software, SPARKY (19.7%), XEASY (16.1%), NMRVIEW (13.7%) and CCPNMR (12.0%), are the most used ones (Figure 1a). Also a great slice of the sector graph belongs to uninformed software (17.1%), which information misses on PDB files and in the correspondent publication, when available. Regarding structure calculation software, CYANA (28.1%), XPLOR-NIH (26.1%), DYANA (a previous version of CYANA, 10.1%) and X-PLOR (a previous version of XPLOR-NIH, 8.5%), are the most used structure calculation software when considering the antimicrobial peptide class of peptides. ARIA represents only 3.3% of this proportion (Figure 1b).

**Figure 1.**
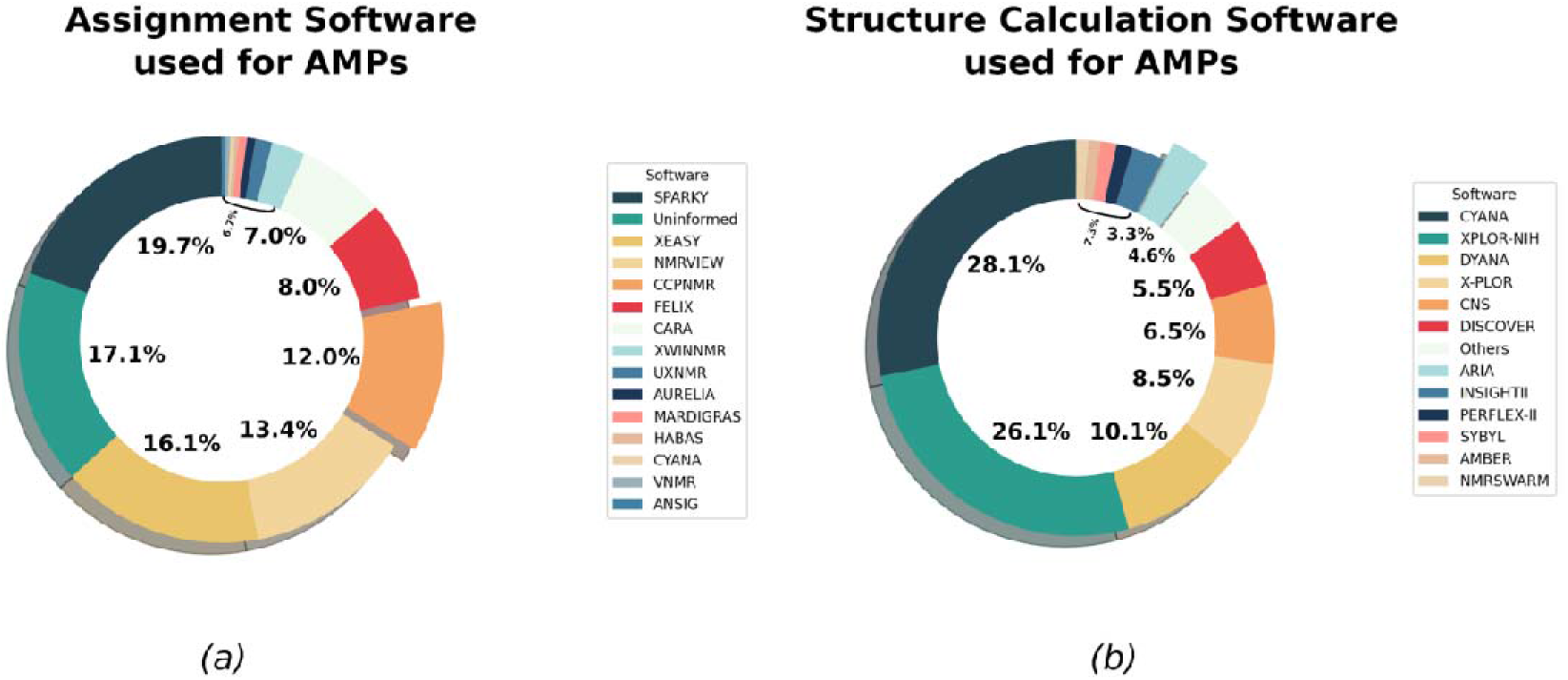
Software used on antimicrobial peptides structure solving. A) Percentage of different assignment software used on PDB deposited AMPs. B) Percentage of different structure calculation software used on PDB deposited AMPs. Detached pie pieces highlight CcpNmr (a) and Aria (b).

To characterize the CcpNmr profile and its impact on PDB deposited structures, the protein structures assigned with CcpNmr were collected from PDB webserver (Table S2). The PDB data allow us to observe that the highest frequency of structure calculation software used with CcpNmr belongs to CYANA (46.7%), with ARIA in the second place (31.2%) (Figure 2a). This result is probably due to the pre-compiled intercommunication between CcpNmr and CYANA/ARIA software. Also, when evaluating the rate of N-terminal acetylated and C-terminal amidated structures in comparison with the other proteins assigned with CcpNmr, we observe an index of 5.1% of 430 proteins, in contrast with XEASY (6.5% of 976 proteins), SPARKY (3.7% of 2100 proteins) and NMRVIEW (2.5% of 2428 proteins) (Figure 2b, Table S3). Even though it is not a low rate, when observing the other tools, the absolute number of proteins of CcpNmr is the lowest. Changing the observation point to ARIA, CcpNmr represents the second software of choice in ARIA absolute numbers registered in PDB database (25%), only behind of NMRVIEW, with 29% (Figure 2c). The scenario changed since CcpNMR came out, and a great burden of the combination CcpNmr/ARIA is observed when CcpNmr Analysis version 2 was released, what made one of its competitor, SPARKY, be relegated from the second to the third place, with 20.5% (Figure 2c, Table S4). In regard to the N-terminal acetylated and C-terminal amidated proteins, the ratio of proteins solved by ARIA is 3.4% (Figure 2d). These results lead us to suggest that, in both ways, CcpNmr and ARIA are one of the main choices for the users of these tools. However, comparing the ratio of N-terminal acetylated and C-terminal amidated proteins to the total proteins assigned in CcpNmr and/or solved by ARIA, along with the relative low ratio of these tools in modified AMPs, we have informations to support the hypothesis that these tools are underused for N-terminal acetylated and C-terminal amidated proteins. A well established protocol for these kind of molecules could improve the usage of CcpNmr and ARIA for N-terminal acetylated and C-terminal amidated proteins.

**Figure 2.**
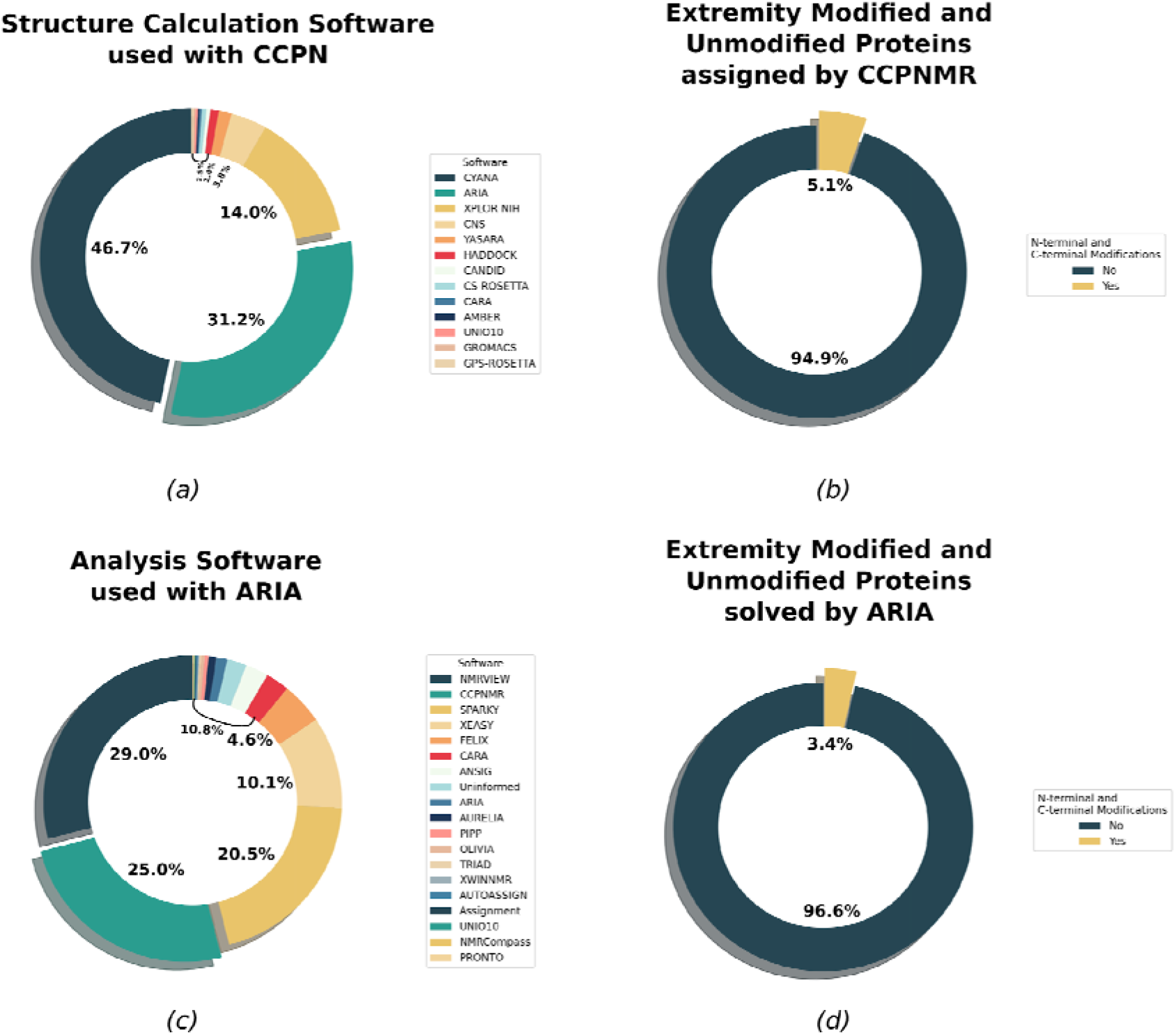
CcpNmr and ARIA pie-chart. All protein structures deposited in PDB which used CcpNmr (a,b) and ARIA (c,d) was accessed. A) The structure calculation software used with proteins assigned on CcpNmr Analysis. B) Representation of N-terminal acetylated and C-terminal amidated structures against non-N-terminal acetylated and non-C-terminal amidated structures. C) The analysis software used with proteins solved by ARIA. D) Representation of N-terminal acetylated and C-terminal amidated structures against non-N-terminal acetylated and non-C-terminal amidated structures. Detached pie pieces represent Aria (a), CcpNmr (c) and modified proteins (b, d).

### 3.2. L-Phenylseptin structures solved with ARIA have similar models with the structures solved with XPLOR-NIH

To verify the implementation of the workflow in Aria2.3, we solved the structure of L-Phenylseptin peptide in DPC micelles. This peptide is found in amphibians with its C-terminal amidated. The NMR spectra assignment was performed by simultaneous analysis of H^1^-H^1^ TOCSY and NOESY, as proposed by Wüthrich. The H◻ and Hβ were used by Dangle tool, present in CcpNmr Analysis, to predict dihedral angles. Except the first phenylalanine, all residue were identified in the experiments and were correlated with at least one more residue (Figure 3). The amide hydrogens allowed us to perform a sequential assignment, which confirms the peptide sequence and provides characteristic correlations on peptide two-dimensional structure. In the NOESY spectra, the most present correlations were NN(i,i+1) and NN(i,i+2). The NN(i,i+1) correlations are present between 3Phe and 4Asp and, after that, 8Asn beyond. The missing residues are complemented with NN(i,i+2) correlations, present in 2Phe with 4Asp and 5Thr with 7Lys. A new NN(i,i+2) correlation is only seen again along 11Lys and 13Ile. A suspect of alpha-helix structure is supported when ◻N(i,i+3) is observed from 2Phe until 16Leu. Also, from 6Leu until 15Ala, there are ◻β(i, i+3) correlations, supported by two ◻N(i, i+4) correlations in the same region, 9Leu with 13Val and 13Val with 17Leu. This results suggest a more strong folded C-terminal portion in comparison with the N-terminal. The alpha region showed 27 short-range NOEs (HN, H◻ − i, i + 1, i+2) and 8 medium range NOEs (HN, H◻ − i, i+3 and i,+4), that, along with short range amide correlations, are characteristic of alpha-helix structures (Fig. 3, table 1).

**Figure 3.**
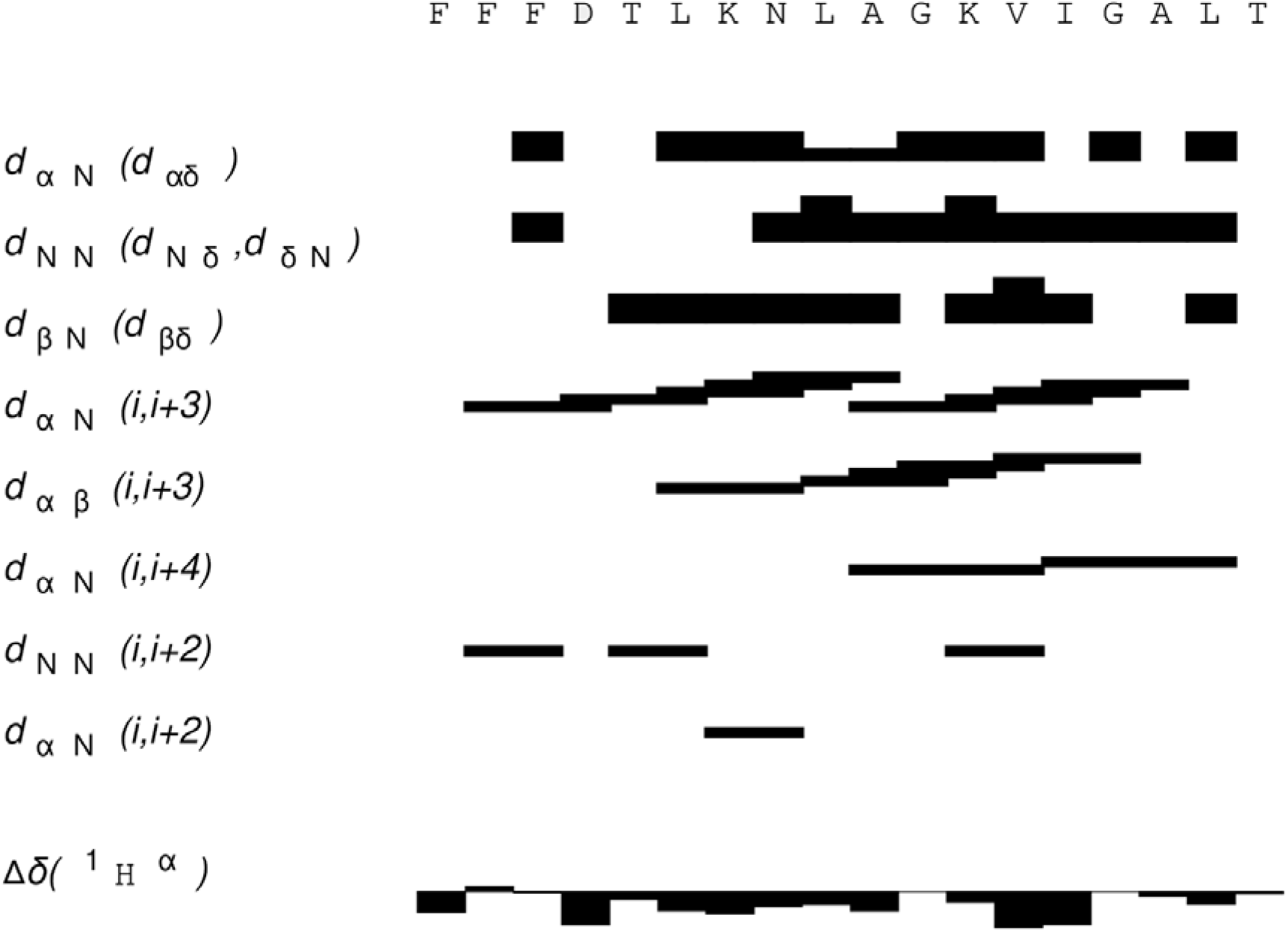
Experimental restraints for L-Phenylseptin. Hydrogen atoms correlations: dαN (hydrogen-alpha and amidic hydrogen), dNN (neighbours amidic hydrogens), dβN (hydrogen-beta and amidic hydrogen), dαN (i,i+3) (hydrogen-alpha in position i and amidic hydrogen in position i+3), dαβ (i, i+3) (hydrogen alpha in position i and hydrogen-beta in position i+3), dαN (i,i+4) (hydrogen-alpha in position i and amidic hydrogen in position i+4), dNN (i,i+2) (amidic hydrogen in position i, amidic hydrogen in position i+2), dαN (i,i+2) (hydrogen-alpha in position i and amidic hydrogen in position i+2), Δδ(^1^H^α^) (hydrogen-alpha secondary shifts). The primary structure is shown in the top. Sequential N-N and α-N indicated by back bars: the thicker the bar, the stronger the connection.

**Table 1.**
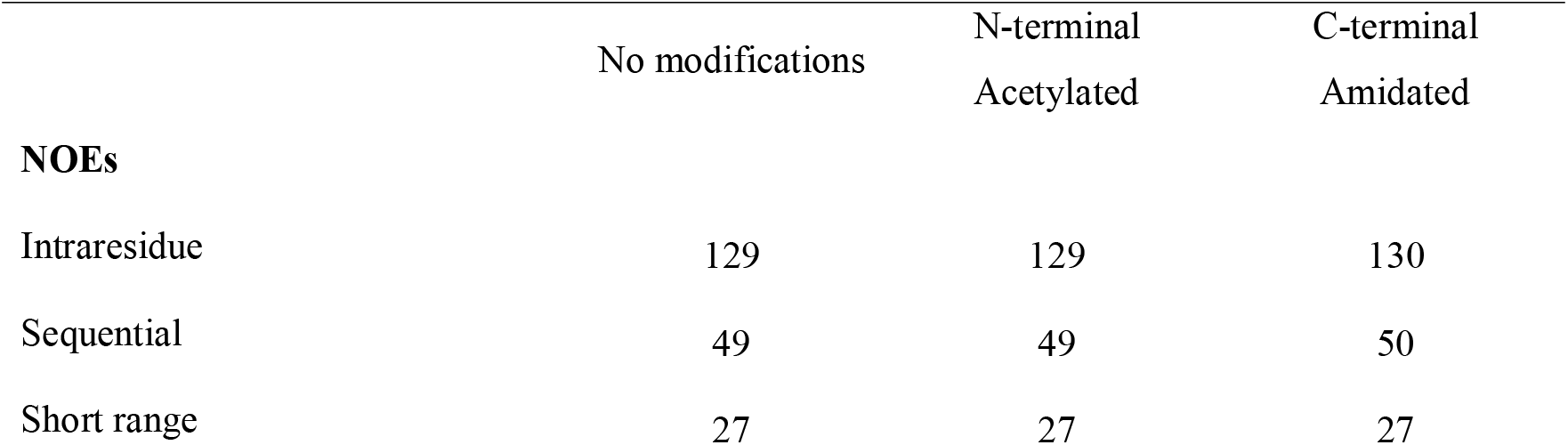
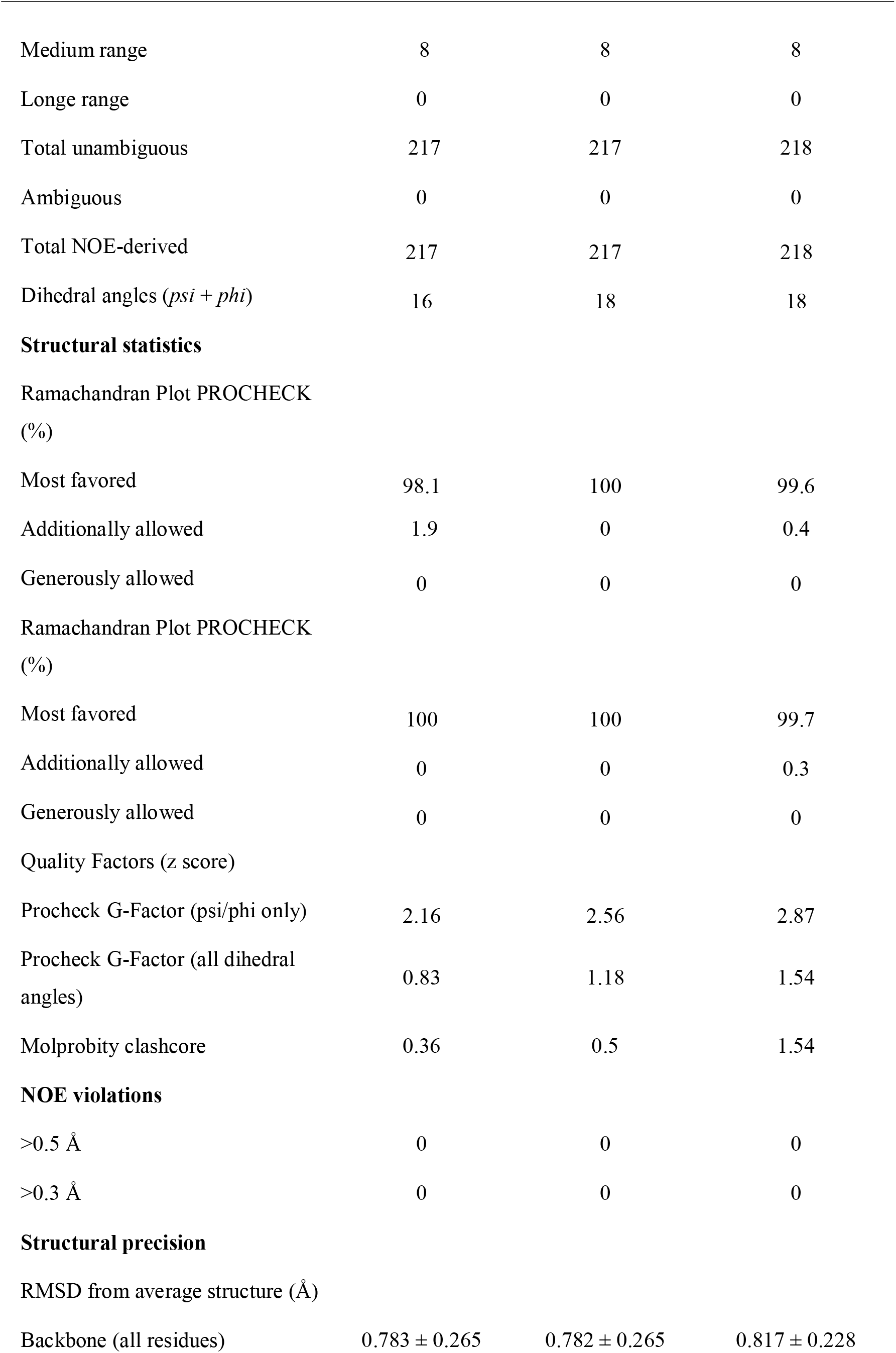
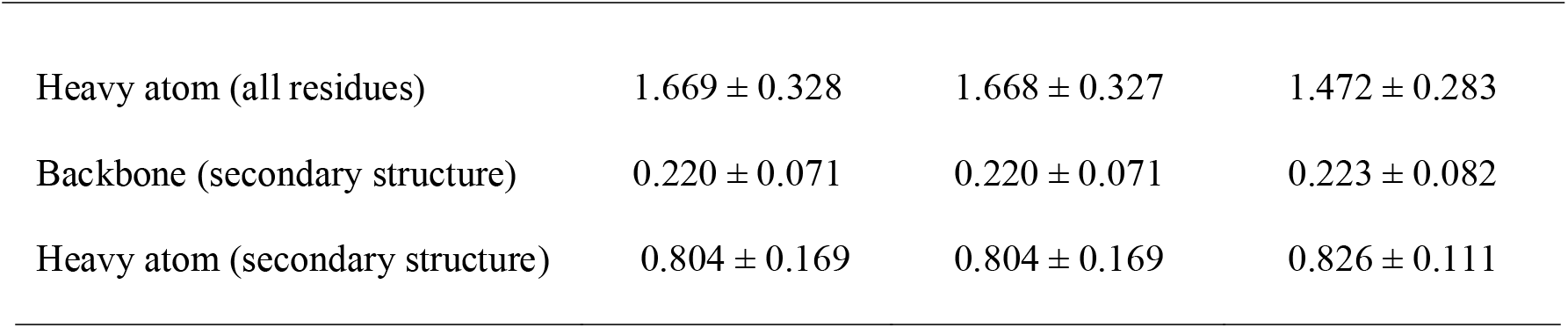
Summary of 20 solution NMR structural statistics Values for the outer shell are given in parentheses.

**Table 2.** Chemical Shift Table of L-Phenylseptin This is a table headnote (style: IUCr table headnote)

|  | H | HA | HB | HG | HD | HE |
| --- | --- | --- | --- | --- | --- | --- |
| 1 Phe | - | 4.25 | 3.12, 2.95 | - | 7.41 | 7.30 |
| 2 Phe | 7.87 | 4.53 | 2.85 | - | 7.33 | 7.15 |
| 3 Phe | 8.50 | 4.50 | 3.06, 2.99 | - | 7.30 | 7.06 |
| 4 Asp | 7.28 | 4.30 | 3.02, 3.01 | - | - | - |
| 5 Thr | 7.78 | 4.23 | 4.05 | 1.26 | - | - |
| 6 Leu | 7.81 | 4.11 | 1.64 | - | 0.89, 0.92 | - |
| 7 Lys | 7.87 | 3.97 | 1.84 | 1.43 | 1.53 | - |
| 8 Asn | 7.89 | 4.53 | 2.94, 2.83 | - | 7.41, 6.77 | - |
| 9 Leu | 8.13 | 4.17 | 1.79, 1.75 | 1.72 | 0.90 | - |
| 10 Ala | 8.52 | 4.03 | 1.46 | - | - | - |
| 11 Gly | 8.02 | 3.82, 3.93 | - | - | - | - |
| 12 Lys | 7.81 | 4.19 | 2.14, 2.00 | 1.53 | 1.63, 1.71 | - |
| 13 Val | 8.15 | 3.68 | 2.22 | 1.04, 0.95 | - | - |
| 14 Ile | 8.66 | 3.76 | 1.91 | 0.92, 1.21,<br>1.74 | 0.83 | - |
| 15 Gly | 8.20 | 3.78, 3.89 | - | - | - | - |
| 16 Ala | 7.94 | 4.24 | 1.57 | - | - | - |
| 17 Leu | 8.30 | 4.24 | 1.92 | 1.58 | 0.87 | - |
| 18 Thr | 7.84 | 4.35 | 4.34 | 1.29 | - | - |

This is a table footnote (style: IUCr table footnote)

Once the NOEs were converted in distance restrictions in CcpNmr, the data was filled in ARIA and the peptide structure solved (Figure 4a). As expected, the molecule is an alpha-helix, with randomized portion among the first two residues and the last residue, due to their lack of correlation, as seen previously (Figure 3). The molecule presents a partial amphipacity, as the hydrophilic residues, 36% of the peptide composition, are in the opposite face to the hydrophobic residue (Figure 4a). The lowest energy structures evidence side-chain rigidity of hydrophobic residues: 3F, 10A, 13V, 14I, 16A, 17L. However, the hydrophilic residues have more free disposition and flexibility (Figure 4c). Since the structure of L-Phenylseptin was already solved in XPLOR-NIH (de Magalhães et al. 2013), we can observe that in macro visualization, no differences were observed between ARIA and X-PLOR-NIH structures (Figure 4b and 4d). The same residues have side-chain rigidity or flexibility.

**Figure 4.**
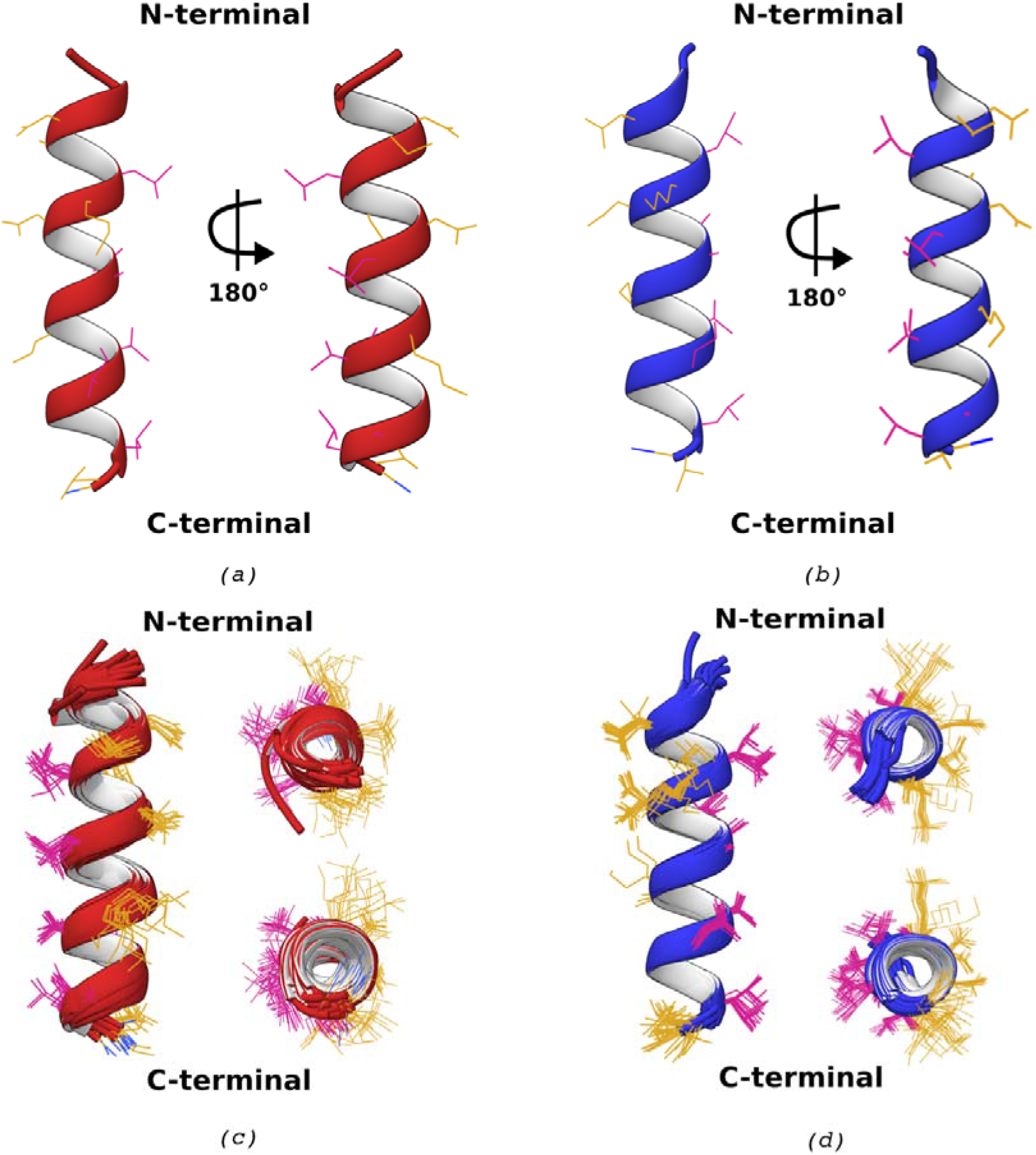
Comparison of D-Phenylseptin structures calculated in ARIA and XPLOR-NIH softwares. A) Peptides structures calculated by Aria2.3, in red. Vertical views from N-terminal until C-terminal. The peptide is represented in two views, after a rotation of 180°. B) Peptides structures calculated by XPLOR-NIH, in blue. Vertical views from N-terminal until C-terminal. The peptide is represented in two views, after a rotation of 180°. Both structures have C-terminal amide modification. C) 20 lowest energy structures of Aria2.3 peptide. Visions: vertical, C-terminal transversal and N-terminal transversal. C) 20 lowest energy structures of Xplor-NIH peptide. Visions: vertical, C-terminal transversal and N-terminal transversal.

The statistical data are present in table 1 and the chemical shift table is present in table S5. The ensemble of the 20 lowest-energy XPLOR-NIH structures showed: a) all residues backbone RMSD of 0.51 ± 0.17; b) all residues heavy atoms RMSD of 1.55 ± 0.4 (de Magalhães et al. 2013). Aria2.3 ensemble of 20 lowest-energy C-terminal amidated structures showed: a) all residues backbone RMSD of 0.81 ± 0.22; b) all residues heavy atoms RMSD of 1.47 ± 0.28. Aria2.3 structures appears to have backbone atoms more disperse and heavy atoms more rigid. However, considering the standard deviations, our analysis suggests that there is no statistic differences between the Xplor-NIH and Aria2.3 structures.

To compare the effect of the termini modifications on the final structures, the L-phenylseptin structures were calculated without N-terminal acetylation and C-terminal amidation. The overall RMSD in the ensemble of 20 lowest energies had no difference between all residues backbone RMSD (0.78 ± 0.26) and all residues heavy atoms RMSD (1.669 ± 0.32 vs 1.668 ± 0.32) (table 1). A slightly difference is observed when no modified structures and N-terminal acetylated structures are compared with C-terminal terminal amidated structures, where all residues backbone RMSD (0.81 ± 0.22) have greater mean value and all residues heavy atoms RMSD (1.47 ± 0.28) have lower mean value. Statistically, we observe that there is no difference between the ensemble of 20-lowest energy structures, however, the lowest structure of the C-terminal amidated peptide seems better folded (Fig 5), as indicated in the tendency of lower heavy atoms RMSD.

**Figure 5.**
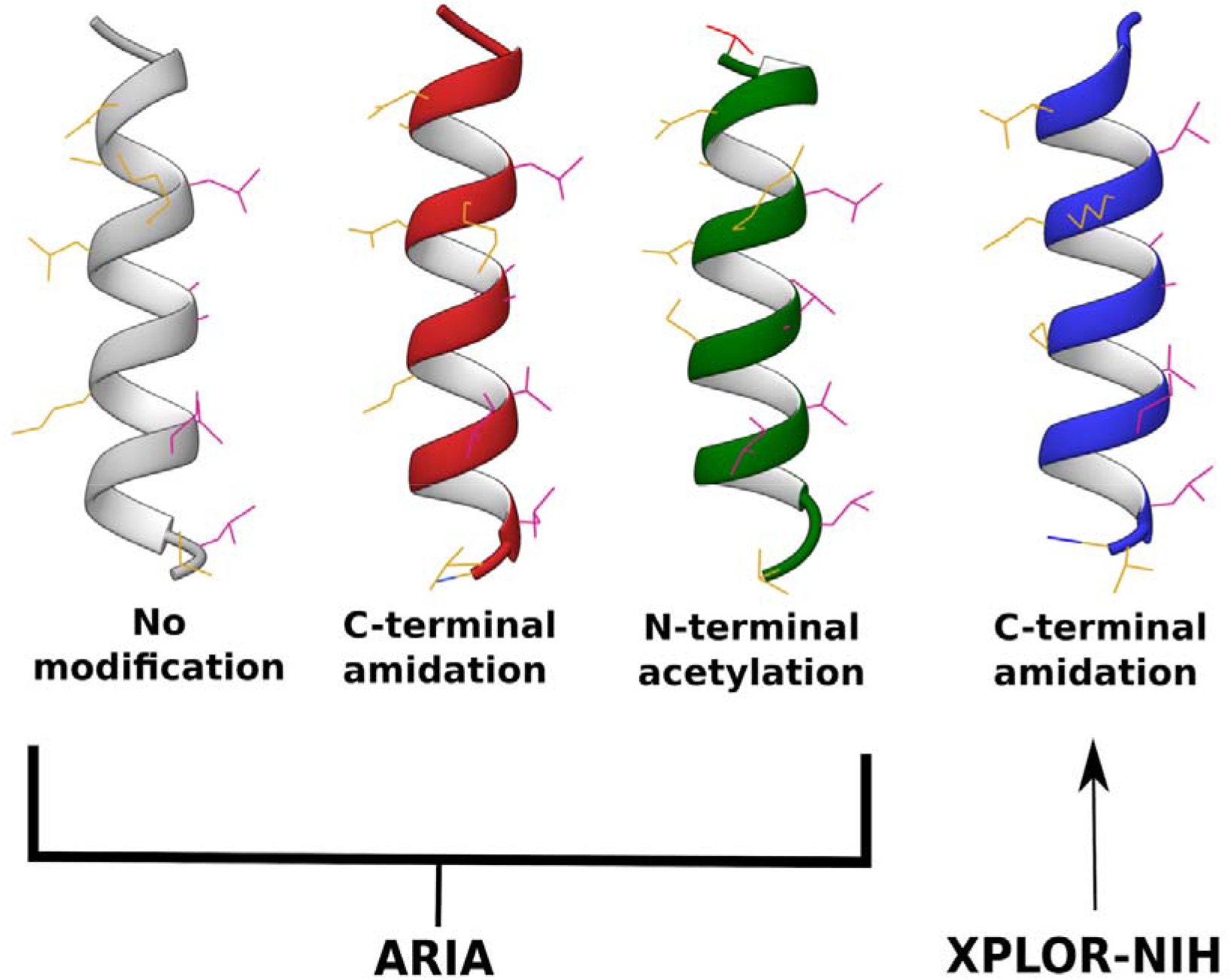
Comparison of peptides C-terminal and N-terminal modification. First structure represents the D-phenylseptin structure without C-terminal or N-terminal modifications. Second structure represents the D-phenylseptin peptide with a C-terminal amidation (dark blue). Third structure represents the peptide calculated with a N-terminal acetylation (red). All three cited structures were calculated using ARIA. Fourth structure contains a C-terminal amidation and was calculated in XPLOR-NIH. All structures represent the lower energy structure between 20 refined in water.

## 4. Conclusions

In this study we improved CcpNmr/ARIA analysis developing a method to implement N-terminal acetylation and C-terminal amidation in proteins for users of CcpNmr Analysis v2.4 and Aria2.3 software. We understand that NMR tools need improvements and we hope to help the scientific academy supporting free aids to the already known software.

## Acknowledgements

BPSO, BSM and MTQM acknowledge grants from CNPq, FAPEMIG and CAPES. This work is a collaboration research project of members of the Rede Mineira de Imunobiológicos (MG) supported by FAPEMIG.

